# wft4galaxy: A Workflow Tester for Galaxy

**DOI:** 10.1101/132001

**Authors:** Marco Enrico Piras, Luca Pireddu, Gianluigi Zanetti

**Author notes:** This work is licensed under a Creative Commons Attribution-NonCommercial-NoDerivatives 4.0 International License (CC BY-NC-ND 4.0).

## Abstract

**Motivation:** Workflow managers for scientific analysis provide a high-level programming platform facilitating standardization, automation, collaboration and access to sophisticated computing resources. The Galaxy workflow manager provides a prime example of this type of platform. As compositions of simpler tools, workflows effectively comprise specialized computer programs implementing often very complex analysis procedures. To date, no simple way exists to automatically test Galaxy workflows and ensure their correctness has appeared in the literature.

**Results:** With `wft4galaxy` we offer a tool to bring automated testing to Galaxy workflows, making it feasible to bring continuous integration to their development and ensuring that defects are detected promptly. `wft4galaxy` can be easily installed as a regular Python program or launched directly as a Docker container – the latter reducing installation effort to a minimum.

**Availability:** `wft4galaxy` is available online at https://github.com/phnmnl/wft4galaxy under the Academic Free License v3.0.

**Supplementary information:** Supplementary information is available at http://wft4galaxy.readthedocs.io.

## 1. INTRODUCTION

Typical bioinformatics analyses involve a number of steps to extract information from various forms of raw data; these analysis procedures are often referred to as *workflows* or *pipelines.* The pattern is so common that a number of *workflow managers* have been created [4] to provide high-level platforms on which to implement these procedures, supporting simpler and more robust implementations than would be reasonably feasible with simple shell scripting. Thus, with the help of workflow managers it becomes practical to implement ever more complex workflows – in fact, workflows with tens of steps are not uncommon. The increase in complexity is accompanied by an increased risk of defects. At best, these will crash and interrupt an analysis procedure; at worst, they will produce subtly wrong results which may only be detected much later. Therefore, given the risks, it seems wise to adopt a mitigation strategy: it is the authors’ opinion the workflow development should be as rigorous as any other kind of software development, especially in light of the growing trend to release and share “standard” workflows. Automated workflow testing then should become an important part of the development process – one which as of yet has not received a lot of attention.

In this work, we present `wft4galaxy`, the WorkFlow Testing tool for the Galaxy data analysis platform [1]. To the best of the authors’ knowledge, `wft4galaxy` is the first published automatic workflow testing tool for Galaxy. `wft4galaxy` works based on the unit testing model: a test case is specified as a set of input datasets and parameters, expected output datasets, and the workflow itself; the workflow is run and the actual and expected outputs are compared. The testing tool uses Galaxy’s RESTful API through the object-orienterd interface of the BioBlend.objects package [5] to automate the entire test execution operation as well as much of the work required to compose the test cases. Of note, our tool is currently used in production within the PhenoMeNal project (http://phenomenal-h2020.eu) to continuously test the workflows integrated in the platform.

## 2. METHODS

The testing model provided by `wft4galaxy` is centered around *test cases.* Each test case defines a workflow and a specific scenario which is to be tested. It contains: the path of the workflow definition file; optionally, the parameters of the various workflow steps; the datasets to be used as workflow inputs; and, finally, expected output datasets. Any number of test cases are collected in a YAML file such as the one shown in Listing 1.

The test definition file is the input for the `wft4galaxy` *test runner,* which automatically executes the entire collection of tests. For each test, the runner connects to an available Galaxy instance provided by the user and then, through the Galaxy API: (1) uploads the workflow; (2) creates a new Galaxy history; (3) uploads all the input datasets; (4) runs the workflow; (5) downloads output datasets. The runner then compares the output to the expected datasets using a comparator function (by default, simple file equality). Finally, all test results are collected and reported.

As an aid to users having to write test definitions, `wft4galaxy` provides a template generator: this tool creates a blank definition and a well-structured directory to contain a test suite.

`wft4galaxy` offers flexibility in the selection of appropriate comparator functions. The default one simply verifies that the files are identical. However, this method is not always appropriate – consider, for instance situations

~~~
 workflows:
  test_case:
   file: “workflow.ga”
   params:
    3:
     “respC”: “gender”
   inputs:
    “DataMatrix”: “input/dataMatrix.tsv”
   expected:
     output:
      file: “expected/variableMetadata.tsv”
     comparator: “comparators.same_row_col”
~~~

**Listing 1**: Example of “Test definition file”

where an analysis may have multiple solutions of comparable quality or cases that are subject to some acceptable degree of round-off error. To handle these cases `wft4galaxy` allows the user to override the default behaviour with customized *comparator functions,* which must be simple Python callables with the signature shown in Listing 2. When specified in a test definition, the custom comparator is automatically loaded and invoked by the `wft4galaxy` framework to decide whether or not the generated output is acceptable for the test. The `wft4galaxy` framework provides a growing package of ready-made comparators (called `wft4galaxy.comparators`), which also includes the default `base_comparator`. Of course, users can also implement their own comparator functions for their tests.

~~~
def my_custom_comparator(generated_file_path, expected_file_path):
 “““ Return True if the two files are “equal”;
  False otherwise. ”””
~~~

**Listing 2**: Signature of a comparator function.

As the individual tests are executed, `wft4galaxy` prints to standard output information about the tests in progress. The format of the output is modelled after the one presented by the Python Unit Test Framework – i.e., for every test case, `wft4galaxy` prints whether it passed or failed. For debugging, more detailed logging can be activated; users can also choose to retain all output files produced by a test run for further analysis and debugging (by default, as soon as the test completes all its datasets and Galaxy histories are deleted).

### Automatic test case generation

The `wft4galaxy` framework further simplifies the creation of workflow test cases through the `wft4galaxy-wizard`, which generates “ready-to-run” workflow test cases from existing Galaxy histories. With the wizard, the steps to create a working test case are reduced to the following. First, the user creates a new history with the required input datasets. Then, the user runs the workflow, after setting any required tool parameters. The workflow should produce a set of new output datasets within the same history. Now, assuming that the workflow has produced correct results, the history can be transformed into a test case by running the `wft4galaxy-wizard`. The wizard will inspect the history to extract and store the underlying workflow (i.e., its ․ga definition file) and all its datasets (both input and output) in a new test directory. The suite definition file is then automatically generated: it will contain a single test case configured to execute the extracted workflow on the input datasets and compare the generated datasets to the outputs of the recorded workflow run.

**Figure 1:**
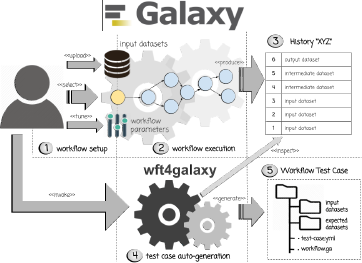
Automatic generation of a test case from a Galaxy history.

### Programmatic Usage

To integrate `wft4galaxy` with third-party tools or for elaborate automation requirements, it can also be used programmatically. Its API is organized around two main packages: `wft4galaxy.core` and `wft4galaxy.wrapper`. The former contains the core logic of the test framework, exposing an Object-Oriented (OO) API for defining and running test cases and test suites programmatically (Listing 3 shows an example of its usage). On the other hand, the latter package contains an OO representation of Galaxy workflows and histories providing a simplified way to inspect inputs, parameters and outputs of tested Galaxy workflows and histories.

~~~
from wft4galaxy.core import
WorkflowTestCase workflow_ filename = “workflow.ga”
inputs = {“ Input Text “ : {“ file “ : “ input “ }}
expected_outputs = {
  “ Out put Text ” : {
   “ file ” : “ expected_outputs ”
 }
}
test_case = WorkflowTest Case (
     base_path, workflow_filename,
inputs, expected_outputs)
test_result = test_case.run(enable_logger=True)
test_result.check_output (“ Out putText “)
~~~

**Listing 3**: Example of programmatic test case definition and execution

### Docker integration

`wft4galaxy` can easily run within a Docker container, completely avoiding any installation hassles. This feature is particularly useful when using continuous integration (CI) services such as Travis CI and Jenkins, where users benefit from not using root privileges for installing new software packages. To simplify the usage of the `wft4galaxy` Docker image, we provide the `wft4galaxy-docker` script, which configures the container execution to use `wft4galaxy` as if it were locally installed. The script can be run standalone, after simply downloading it from the `wft4galaxy` GitHub repository.

## 3. CONCLUSION

`wft4galaxy` is a tool to simplify and automate workflow tests. It supports the adoption of “unit testing” and continuous integration into the workflow development and maintenance process. Its native support for Docker enables easy integration with specialized CI systems, such as Jenkins. Indeed, within the PhenoMeNal project, Jenkins with `wft4galaxy` are used to test complex workflows such as the ones described by De Atauri et al. [3]. Although in its current version `wft4galaxy` is tied to the Galaxy platform, in the future we would like to investigate the feasibility of extending it to work with other workflow management systems and, in particular, implementations of the Common Workflow Language [2].

## Acknowledgements

The authors would like to thank the fellow members of the PhenoMeNal team for their valuable feedback. This work was partially supported by the European Commission’s Horizon2020 programme under the PhenoMeNal project (654241) and by the Region of Sardinia under project ABLE.

## References

[1] E. Afgan, D. Baker, M. van den Beek, D. Blankenberg, D. Bouvier, M. Cech, J. Chilton, D. Clements, N. Coraor, C. Eberhard, B. Grüning, A. Guerler, J. Hillman-Jackson, G. Von Kuster, E. Rasche, N. Soranzo, N. Turaga, J. Taylor, A. Nekrutenko, and J. Goecks. The Galaxy platform for accessible, reproducible and collaborative biomedical analyses: 2016 update. Nucleic Acids Research, 44(May):gkw343, 2016. ISSN 0305–1048. doi:10.1093/nar/gkw343.

[2] P. Amstutz, M. R. Crusoe, N. Tijanić, B. Chapman, J. Chilton, M. Heuer, A. Kartashov, D. Leehr, H. Ménager, M. Nedeljkovich, M. Scales, S. Soiland-Reyes, and L. Stojanovic. Common Workflow Language, v1.0. 1 2016. doi:10.6084/M9.FIGSHARE.3115156.V2.

[3] P. De Atauri, P. Moreno, V. Selivanov, C. Foguet, S. Marin, S. Neumann, and M. Cascante. Workflows For Fluxomics In The Framework Of Phenomenal Project. 1 2016. doi:10.5281/ZENODO.154584.

[4] J. Leipzig. A review of bioinformatic pipeline frameworks. Briefings in Bioinformatics, (January):bbw020, 3 2016. ISSN 1467–5463. doi:10.1093/bib/bbw020.

[5] S. Leo, L. Pireddu, G. Cuccuru, L. Lianas, N. Soranzo, E. Afgan, and G. Zanetti. BioBlend.objects: Metacomputing with galaxy. Bioinformatics, 30(19):2816–2817, 2014. ISSN 14602059. doi:10.1093/bioinformatics/btu386.

